# A tightly regulated auxin signaling landscape is required for spatial accommodation of lateral roots in *Arabidopsis*

**DOI:** 10.1101/2025.01.07.631685

**Authors:** Thái X. Bùi, Vinay Shekhar, Sophie Marc-Martin, Kevin Bellande, Joop E.M. Vermeer

## Abstract

In *Arabidopsis thaliana*, lateral root (LR) development requires spatial accommodation responses in overlying endodermal cells. This includes loss of cell volume whilst maintaining membrane integrity to allow the expansion of the underlying LR primordia (LRPs). These accommodation responses are regulated by auxin-mediated signaling, specifically through Aux/IAA proteins, involving *IAA3/SHY2*. Plants that express a stabilized version of SHY2, shy2-2, in differentiated endodermal cells, *CASP1_pro_::shy2-2* plants, fail to make LRs. Exogenous treatment with 1- naphthaleneacetic acid (NAA) was reported to partially restore LR formation in this spatial accommodation mutant. Using treatments with auxins with different transport properties, such as NAA, indole-3-acetic acid (IAA), and 2,4-dichlorophenoxyacetic acid (2,4-D), we assessed the ability of each auxin to rescue LR formation in *CASP1_pro_::shy2-2* roots. This revealed that IAA is the most effective in partially restoring LR development, NAA is effective in inducing LRPs but cannot maintain their canonical phenotype, whereas 2,4-D induces non-controlled cell divisions. In addition, we show that in *CASP1_pro_::shy2-*2 roots, *AUX1* appears to be repressed in the zone where oscillation of the auxin response have been described. Our study advances the understanding of auxin-regulated spatial accommodation mechanisms during LRP formation and highlights the complex interplay of auxin transport and signaling in bypassing the endodermal constraints.

## INTRODUCTION

The root system plays an important role in plant growth and development. The regulated formation of LRs gives rise to a belowground network, which is a major determinant for nutrition uptake, anchorage, storage and rhizosphere interactions (Péret *et al*., 2009; Schäfer *et al*., 2022; Gifford *et al*., 2024). In addition, LR development has been shown to be able to perceive and react to environmental cues to contribute to plant adaptation (Perotti *et al*., 2020; Schäfer *et al*., 2022). In *Arabidopsis*, LRs initiate from a set of the xylem pole pericycle (XPP) cells, called LR founder cells. To emerge, the newly formed organ must traverse three overlying cell layers: the endodermis, cortex and epidermis, respectively. Multiple aspects of LR development, from positioning to initiation, outgrowth and emergence, have been known to be regulated by the phytohormone auxin and its signaling pathways (Lavenus *et al*., 2013; Du & Scheres, 2018). One branch of auxin signaling is marked with the binding of IAA, the plant endogenous auxin, to the co-receptor complex consisting of the F-box TRANSPORT INHIBITOR RESPONSE1 (TIR1) protein and Aux/IAA proteins, leading to the ubiquitination and degradation of Aux/IAAs by the 26S proteasome (Zenser *et al*., 2001; Gray *et al*., 2003; Dharmasiri *et al*., 2005). Since Aux/IAAs are repressors of AUXIN RESPONSE FACTORS (ARFs), transcription factors that regulate auxin- dependent gene expression, their degradation releases the inhibition of ARFs, triggering auxin responses (Guilfoyle & Hagen, 2007; Goh *et al*., 2012; Cancé *et al*., 2022).

In *Arabidopsis*, LR formation requires Aux/IAA-mediated signaling in the XPP as well as in the neighboring endodermis (Vanneste *et al*., 2005; De Smet *et al*., 2007, 2010; De Rybel *et al*., 2010; Vermeer *et al*., 2014; Santos Teixeira & ten Tusscher, 2019). In the LR founder cells, Aux/IAA- mediated signaling was shown to regulate the remodeling of the cytoskeleton to promote asymmetric cell expansion and nuclear migration to execute the formative divisions resulting in a stage I LRP (Vilches Barro *et al*., 2019). Concomitantly, Aux/IAA mediated responses in the endodermis include remodeling of the microtubule organization as well as a regulated loss of cell volume (Stoeckle *et al*., 2022). These responses in the endodermis were coined spatial accommodation responses, as these cellular adjustments accommodate the expansion growth of the new LR (Vermeer *et al*., 2014; Stoeckle *et al*., 2018, 2022). The importance of Aux/IAA mediated signaling in the endodermis was revealed via the expression of a stabilized allele of *SHORT HYPOCOTYL2* (*SHY2*) / *IAA3*, a dominant repressor of auxin-mediated gene expression, in differentiated endodermal cells (Vermeer *et al*., 2014). In this mutant, *CASP1_pro_::shy2-2*, LR formation was completely abolished. It was proposed that these auxin-mediated responses are required to trigger mechanical feedback from the endodermis to sustain the expansion growth of the new LR (Vermeer *et al*., 2014; Vermeer & Geldner, 2015). Interestingly, the application of NAA was shown to be able to partially restore LR formation in *CASP1_pro_::shy2-2* (Vermeer *et al*., 2014). However, most of the resulting LRs were often malformed and their emergence was strongly impaired, but the reason why NAA only partially rescued the *CASP1_pro_::shy2-2* LR phenotype remains unknown (Vermeer *et al*., 2014).

Thus, we treated the *CASP1_pro_::shy2-2* mutant with different auxin analogues (IAA, NAA and 2,4- D) to explore their potential in rescuing its LRP formation. Their unique stability levels and transport requirements (Fig. S1) allowed us to characterize auxin responses in *CASP1_pro_::shy2-2*. Our results revealed that IAA is the most potent of the tested auxins to restore LRP formation, whereas NAA and 2,4-D were less capable. In addition, the exogeneous auxin treatments could readily overcome the block of cell divisions in the XPP, but appeared to be less efficient in restoring the Aux/IAA-mediated responses in the endodermis. Our results suggest that the block in auxin mediated responses in the endodermis of *CASP1_pro_::shy2-2* is so strong that even with exogenous auxin treatments the recovery of auxin signaling is limited. Furthermore, we also investigated whether auxin signaling, cytokinin signaling and the expression of some of the major auxin transporters known to be involved in LR formation, like PIN-FORMED1 (PIN1) and AUXIN RESISTANT 1 (AUX1) (Bennett *et al*., 1996; Benková *et al*., 2003), were modified in *CASP1_pro_::shy2-2*. The observed delayed expression of AUX1 in the XPP of *CASP1_pro_::shy2-2*, roots suggested that auxin-mediated communication between the XPP and endodermis could be required for AUX1 accumulation in the XPP to potentiate LR formation.

## MATERIALS AND METHODS

### Plant materials

We used Columbia-0 (Col-0), *CASP1_pro_::shy2-2* (Vermeer *et al*., 2014) and their reporters, including *UBQ10_pro_::EYFP:NPSN12* (Wave131Y; Geldner *et al*., 2009), *DR5::NLS-3xVENUS* (Heisler *et al*., 2005), *TCSn::GFP* (Zürcher *et al*., 2013), *AUX1:mVenus* (Swarup *et al*., 2004) and *PIN1:GFP* (Benková *et al*., 2003). All transgenic lines were in Columbia-0 (Col-0) background and introgressed into *CASP1_pro_::shy2-2*.

### Culture conditions

Seeds were sterilized in chlorine gas and sown on half-strength Murashige and Skoog (MS) medium with 2.4 g L^-1^ MS basal salt (Duchefa Biochemie) and 0.8% (m/v) plant agar (Duchefa Biochemie). After 48 hours of stratification in the dark at 8°C, they were transferred into the growth chamber (Aralab, 22°C, continuous light).

### Auxin treatment assay

IAA, NAA or 2,4-D were dissolved in dimethyl sulfoxide (DMSO), and each added into the half-strength MS medium at the concentration of 0.1, 1 or 10 µM. DMSO was factored into the medium for the control condition accordingly. 5-day old *Arabidopsis* seedlings were transferred into auxin treatment plates and kept in the growth chamber (Aralab, 22°C, continuous light) for up to 48 or 72 hours prior to harvest.

### Root clearing and phenotyping

Harvested seedlings of Col-0 and *CASP1_pro_::shy2-2* were cleared by immersion in 20% (v/v) methanol/ 4% (v/v) HCl solution at 57°C for 20 minutes, then in 7% (w/v) NaOH/ 60% (v/v) ethanol solution at room temperature with gentle agitation for 15 minutes, followed by a three-step rehydration using 40%, 20% and 10% (v/v) ethanol solutions. Cleared samples were stored in 50% (v/v) glycerol at 4°C and mounted onto glass slides for phenotyping. LRP number and developmental stages of Col-0 and *CASP1_pro_::shy2-2* plants were determined on cleared samples (*n* ≥ 18 seedlings per condition, in three replicates) and imaged using a Leica DM4 B microscope operating in differential interference contrast (DIC) mode. LRP width and height were measured using LAS X software (*n* = 20 LRPs per stage (stage I – IV, prior to LRP crossing the endodermis), per condition, in three replicates). Statistical analysis was done using RStudio Desktop software (open-source edition, Posit PBC).

### Multiphoton microscopy and image analysis

LRP morphology, auxin and cytokinin signaling, and auxin transport were investigated by using a Leica SP8 multiphoton confocal microscope with 4Tune detector operating in counting mode with 960nm excitation wavelengths and 510-600nm detection window for green/yellow fluorescent proteins running Leica LAS X software (Stoeckle *et al*., 2022; de Jesus Vieira Teixeira *et al*., 2024). Seedlings of the plasma membrane reporter lines (*UBQ10_pro_::EYFP:NPSN12*), were imaged live. Seedlings of the other reporters (*DR5::NLS-3xVENUS*, *TCSn::GFP*, *AUX1:VENUS* and *PIN1:GFP*) were fixed and cleared following the ClearSee method (Kurihara *et al*., 2015) prior to being imaged. All obtained images were processed and analyzed using Fiji software (Schindelin *et al*., 2012).

## RESULTS

### Exogeneous IAA, NAA and 2,4-D addition can only partially restore LR formation in *CASP1_pro_::shy2-2* roots

The inability of *CASP1_pro_::shy2-2* plants to form LRs is due to a dominant repression of Aux/IAA-mediated auxin signaling in differentiated endodermal cells (Vermeer *et al*., 2014). Here we applied treatment with different auxins to investigate their capacity to rescue the LR phenotype in *CASP1_pro_::shy2-2* roots. Specifically, 5-day old Col-0 (wild type) and *CASP1_pro_::shy2-2* seedlings were treated with IAA, NAA and 2,4-D at different concentrations (Fig. 1). We observed that untreated wild type seedlings exhibit emerged LRs with wide spacing and asymmetrical distributing patterns, as described by others, along the primary root. However, none of the treatments with the tested auxins could evoke a similar LR phenotype in *CASP1_pro_::shy2-2* (Fig. 1). Although 2,4-D massively induces cell divisions in the XPP in both backgrounds, the treatment did not result in the formation of LRs with the canonical dome-shaped morphology (Fig. 1A & D). Even in wild type, 10 μM 2,4-D triggered an upsurge of cell proliferation in the pericycle without maintaining proper spacing between successive LRPs/LRs to a point where individual LRPs could no longer be distinguished (Fig. 1A & D). Occasionally, we observed LRPs traversing the endodermis or emerging from the cell proliferation zone along the primary root (Fig. 1A, D, G, J & S2C). In *CASP1_pro_::shy2-2*, 10 μM 2,4-D application also resulted in a cell proliferation zone spanning along the primary root without separate LRPs, but this zone appeared flat compared to wild type (Fig. 1A & D). Prolonged treatment could trigger more cell divisions, but *CASP1_pro_::shy2-2* plants rarely formed distinct LRPs and the dividing cells struggled to traverse the endodermis compared to wild type (Fig. 1G, J & S2C).

**Figure 1.**
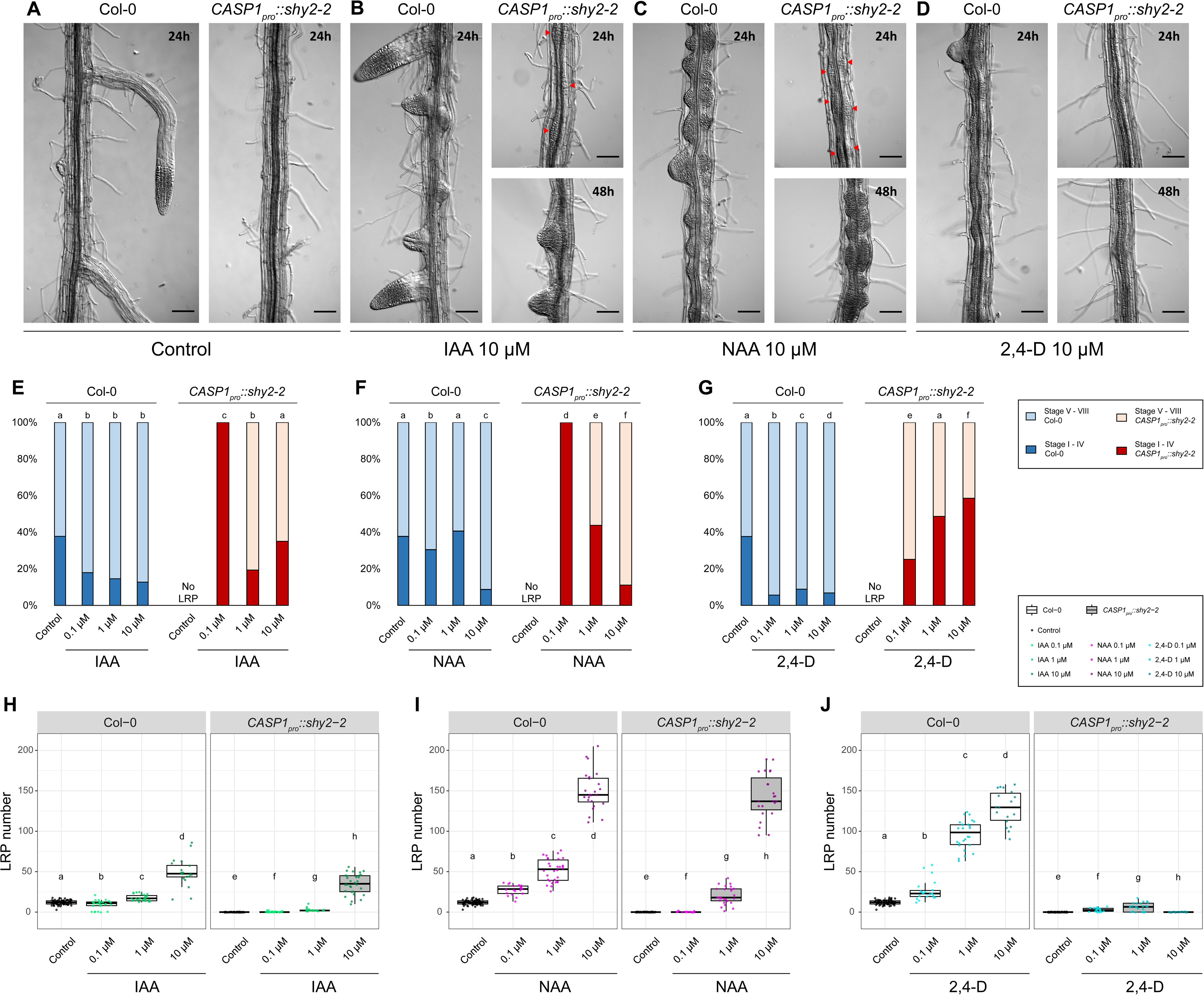
IAA and NAA induce LR formation in *CASP1_pro_::shy2-2* with a delay, whereas 2,4- D is less efficient. **A-D**. LRP induction after 24-hour and 48-hour 10 μM auxin treatments in Col-0 and *CASP1_pro_::shy2-2*. Red arrowheads indicate LRPs induced by IAA and NAA after 24-hour treatment. Scale bar = 100 μm. **E-G**. Proportion of LRP stages before traversing the endodermis (stage I - IV) in Col-0 (dark blue) and *CASP1_pro_::shy2-2* (dark red) and after (stage V - VIII) in Col-0 (light blue) and *CASP1_pro_::shy2-2* (light red), induced by IAA (**E**), NAA (**F**) and 2,4-D (**G**). Different letters indicate Pearson’s χ^2^ test significant difference (p-value < 0.05). **H-J**. LRP number of Col-0 and *CASP1_pro_::shy2-2* after 48 hours of IAA (**H**, green), NAA (**I**, magenta) and 2,4-D (**J**, cyan) treatments. Different letters indicate Wilcoxon rank-sum test significant difference (p-value < 0.05). 5-day old seedlings of Col-0 and *CASP1_pro_::shy2-2* were treated with IAA, NAA and 2,4-D at 0.1, 1 or 10 μM for up to 48 hours (*n* ≥ 18 plants per condition, in three replicates).

NAA treatment also resulted in a strong induction of cell divisions in the XPP, but unlike 2,4-D, it did not completely disrupt the organization of LRPs in *CASP1_pro_::shy2-2*. NAA treatments significantly increased the number of LRPs in both backgrounds (Fig. 1I). Indeed, LRPs are distinguishable, though very densely distributed on a cell proliferation zone along the primary root, with reduced spacing distance between each two successive LRPs (Fig. 1A & C). Interestingly, in *CASP1_pro_::shy2-2*, none of the formed LRPs managed to traverse the endodermis after 48 hours of 0.1 μM NAA treatment. However, raising the NAA concentration led to a higher percentage of LRPs successfully traversing the endodermis and emerging (Fig. 1F & S2B). Nonetheless, independent of the used concentration of NAA, we observed inconsistent LRP morphology and a strong a delay in LR emergence, most likely due to the problems with traversing the endodermis (Fig 1A & C).

IAA treatment was the most potent in reverting the LR phenotype of *CASP1_pro_::shy2-2* plants. Treatment with 10 µM IAA did not trigger extensive cell divisions in the XPP, nor did it produce a cell proliferation zone along the primary root comparable to that observed in roots treated with NAA or 2,4-D. In fact, LRP shape, spacing distance and distribution in *CASP1_pro_::shy2-2* treated with 10 μM IAA were even more resemblant to wild type (Fig. 1A & B). The LRP count of IAA- and NAA-treated *CASP1_pro_::shy2-2* plants both increased significantly, albeit the IAA-treated displayed a much smaller margin of increase (Fig. 1H & I). In *CASP1_pro_::shy2-2*, 0.1 μM IAA treatment induced very few LRPs, all of which appeared to be arrested in their early stages of development, whereas treatments with 1 μM and 10 μM IAA resulted in higher percentage of LRPs traversing the endodermis and emerging (Fig. 1E & S2A). Additionally, LR emergence in *CASP1_pro_::shy2-2* recovered by IAA treatment was also delayed compared to the wild type (Fig. 1A, B & S2A).

Overall, by assessing the responsiveness of *CASP1_pro_::shy2-2* to different auxins, we have shown that the LR phenotype in *CASP1_pro_::shy2-2* could only be partially recovered by exogenous auxin treatments, with IAA being the most efficient and 2,4-D the least. Furthermore, the observed delay in LR emergence in *CASP1_pro_::shy2-2* despite treatment with moderate to high concentrations of auxins (Fig. 1A-D & S2), suggests that the spatial accommodation responses in the endodermis are not easily restored by treating them with exogenous auxins.

### Auxin treatment fails to restore stereotypical LR morphology in *CASP1_pro_::shy2-2* roots

To analyze more directly the effect of a resistant endodermis on restricting LR formation in *CASP1_pro_::shy2-2*, we crossed the *UBQ10_pro_::EYFP:NPSN12* plasma membrane reporter (Geldner *et al*., 2009) into the *CASP1_pro_::shy2-2* background (Fig. 2A). Treating seedlings with the different auxins all resulted in a loss of the canonical dome shape of LRPs until stage IV. In *CASP1_pro_::shy2-2*, the LRPs mostly remained flattened against the endodermis compared to wild type (Fig. 2A). The only exception was seen in IAA-treated *CASP1_pro_::shy2-2* plants, in which we observed dome-shaped LRPs, though slightly flatter compared to wild type (Fig. 2A). In addition, we noticed that stage II and IV primordia in *CASP1_pro_::shy2-2* were wider than in Col-0 when treated with IAA or NAA (Fig. 2A). Regarding 2,4-D treatment, the cell proliferation zone in *CASP1_pro_::shy2-2* was clearly consistent with our previous observations and resulted in a completely flat proliferative zone beneath the endodermis in which we could not observe individual LRs (Fig. 1A, D and 2A).

**Figure 2.**
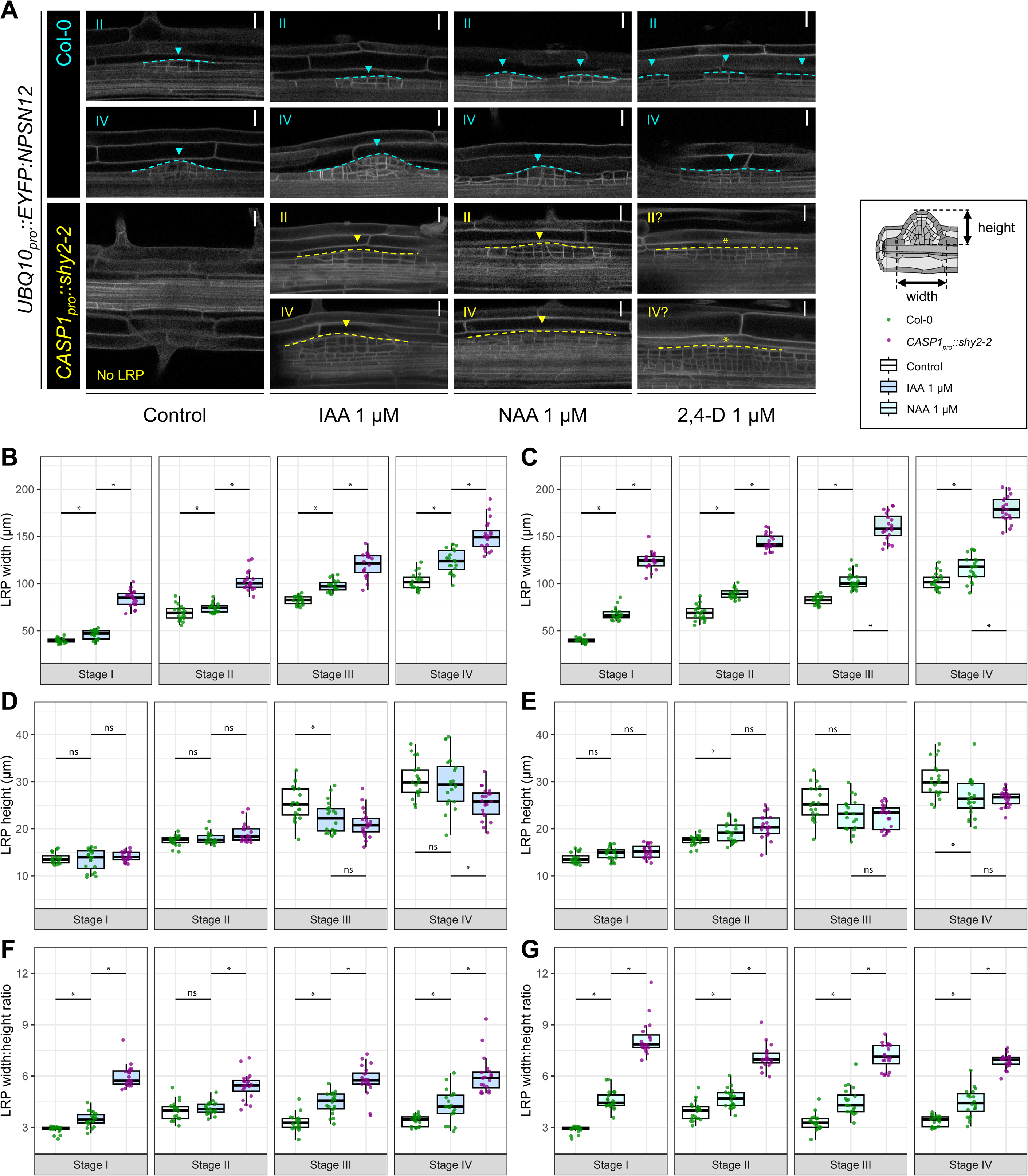
Auxin treatments affect LRP morphology in *CASP1_pro_::shy2-2*. **A**. 2,4-D induces uncontrolled cell divisions without maintaining proper LRP morphology, whereas IAA- and NAA- mediated LRPs in *CASP1_pro_::shy2-2* (yellow) are deformed compared to wild type (cyan). LRP morphology is visualized by the plasma membrane marker *UBQ10_pro_::EYFP:NPSN12* (grey) in Col-0 and *CASP1_pro_::shy2-2* backgrounds. 5-day old seedlings of Col-0 and *CASP1_pro_::shy2-2* were treated with IAA, NAA and 2,4-D at 1 μM for 48 hours. Roman numerals, dashed lines and arrowheads of the same colors indicate LRP stages and positions in Col-0 (cyan) and *CASP1_pro_::shy2-2* (yellow). Asterisks indicate uncontrolled cell division. Scale bar = 20 μm; min. 5 images per observation, 3 independent experiments. **B-G**. IAA and NAA treatments result in widened and flattened LRPs in *CASP1_pro_::shy2-2*. LRP dimensions (width (**B**, **C**), height (**D**, **E**) and ratio between LRP width and height (**F**, **G**)) through every stage before they cross the endodermis (stage I - IV) in Col-0 (green) and *CASP1_pro_::shy2-2* (magenta) were quantified on 20 LRPs per stage, per condition, in three replicates. ANOVA post hoc Student’s *t*-test significant difference is represented by asterisks (p-value ≤ 0.05) or “ns” (non-significant difference, p-value > 0.05).

We analyzed the auxin-induced LRP morphology by quantifying the variation in the morphology of stage I to IV LRPs in both backgrounds after auxin treatment (*CASP1_pro_::shy2-2* plants do not make any LRs without addition of auxin). Particularly, we measured the dimensions of the auxin-induced LRPs prior to stage V, before they have traversed the endodermis. The quantification of LRP dimensions of up-to-stage-IV demonstrates a significant increase in LRP width in *CASP1_pro_::shy2-2* compared to Col-0 whether they are treated with 1 μM IAA or NAA (Fig. 2B & C). Moreover, the ratio between LRP width and height is significantly higher in *CASP1_pro_::shy2-2* compared to wild type with 1 μM IAA and NAA treatments (Fig. 2F & G), confirming that auxin treatments result in a widening and flattening of LRP in *CASP1_pro_::shy2-2*. On the other hand, LRP height in *CASP1_pro_::shy2-2* roots after IAA and NAA treatments either was not significantly affected or was lower (stage-IV; IAA-induced LRPs) compared to wild type (Fig. 2D & E). This suggests that although exogenous auxin treatments can induce cell division and presumably LRP initiation from the XPP, the lack of spatial accommodation responses of the overlying endodermis in *CASP1_pro_::shy2-2* is not easily overcome by auxin treatment and appears to restrain LRP outgrowth, resulting in wider and flatter LRP with a delayed emergence.

### Transport properties of auxins affect induction of auxin-mediated transcriptional responses in *CASP1_pro_::shy2-2*

Aux/IAA-mediated signaling plays a key role throughout LR development. Due to the strong impairment in endodermal Aux/IAA-mediated signaling, *CASP1_pro_::shy2-2* endodermal cells are not responsive to the initiation of expansion growth in the XPP and this results in a complete absence of LRs (Vermeer *et al*., 2014). Since we showed that treatment with auxins with different transport properties leads to various levels of reversion of the LR phenotype in *CASP1_pro_::shy2-2*, we tested if there was a correlation between the level of rescue with the expression of the auxin response marker *DR5::NLS-3xVENUS* (Heisler *et al*., 2005; De Smet *et al*., 2007; Moreno-Risueno *et al*., 2010) (Fig. 3A). Overall, auxin treatments induced *DR5::NLS-3xVENUS* in *CASP1_pro_::shy2- 2* roots, but rather weak compared to wild type roots (Fig. 3A). IAA and NAA treatments resulted in an increased *DR5::NLS-3xVENUS* fluorescence in the *CASP1_pro_::shy2-2* root apical meristem, but slightly lower than Col-0, whereas the effect of 2,4-D on *DR5::NLS-3xVENUS* fluorescence was remarkably low. Aside from that, the induction of *DR5::NLS-3xVENUS* in *CASP1_pro_::shy2-2* roots was more pronounced in the LR formation zone compared to the LR branching zone, independent of the type of auxin used. However, we observed that NAA treatment affected a notably larger area in which *DR5::NLS-3xVENUS* fluorescence accumulated compared to IAA and 2,4-D (Fig. 3A). Nevertheless, the auxin-induced increase in *DR5::NLS-3xVENUS* is mostly restricted within the vasculature of *CASP1_pro_::shy2-2* (Fig. 3A), suggesting that auxin treatments can modify auxin signaling in *CASP1_pro_::shy2-2*, but cannot efficiently bypass the endodermal blockage. In addition, we also observed that the induction of *DR5::NLS-3xVENUS* fluorescence in the cortex and epidermis of *CASP1_pro_::shy2-2* roots was weaker compared to wild type (Fig. 3A). This suggests that the block in Aux/IAA-mediated signaling in the endodermis also has a non-cell autonomous effect on the outer cell layers in addition to the effect on the XPP.

**Figure 3.**
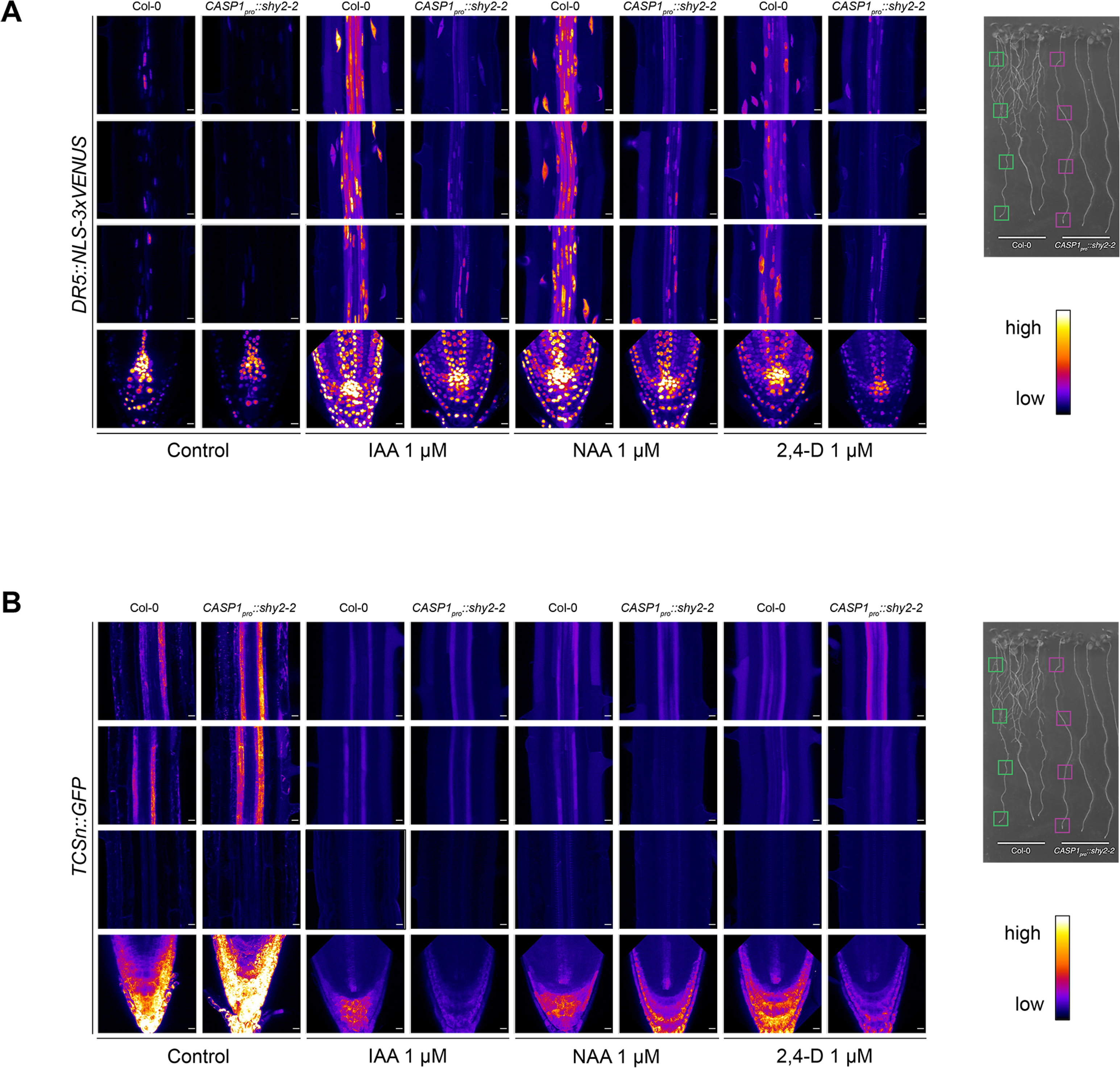
Mapping auxin and cytokinin responses after IAA, NAA or 2,4-D treatment in *CASP1_pro_::shy2-2*. **A**. Auxin signaling outputs are restricted in the vasculature and down-regulated in *CASP1_pro_::shy2-2*. **B**. Cytokinin signaling outputs seem unaffected in *CASP1_pro_::shy2-2*. 5-day old seedlings of reporter lines for auxin signaling, *DR5::NLS-3xVENUS* (**A**), and of cytokinin signaling, *TCSn::GFP* (**B**), with Col-0 and *CASP1_pro_::shy2-2* backgrounds, treated with IAA, NAA and 2,4-D at 1 μM for 4 hours, were fixed and cleared with ClearSee, and imaged using multiphoton microscope. Relative positions of observation along the primary root are marked with green (Col-0) and magenta (*CASP1_pro_::shy2-2*) boxes. Scale bar = 10 μm; min. 5 images per condition, in 3 independent experiments.

### *CASP1_pro_::shy2-2* roots display altered levels of the cytokinin response marker TCSn::GFP

The phytohormone cytokinin acts antagonistically in concert with auxins to co-regulate multiple plant biological processes including LR development (Laplaze *et al*., 2007; Bielach *et al*., 2012; Marhavý *et al*., 2014; Kieber & Schaller, 2018; Nenadić & Vermeer, 2021). Since SHY2 has been linked to the regulation of cytokinin responses (Ioio *et al*., 2008), we mapped cytokinin signaling outputs in *CASP1_pro_::shy2-2* using the *TCSn::GFP* reporter (Zürcher *et al*., 2013). Under control conditions, we observed a slight increase in TCSn::GFP signal in the root tip and the pericycle of the differentiated part of roots of *CASP1_pro_::shy2-2* plants compared to wild type roots (Fig. 3B). Treatment with the different auxins repressed cytokinin signaling both in Col-0 and *CASP1_pro_::shy2-2* compared to control conditions (Fig. 3B). Stronger suppression of TCSn::GFP was observed towards the LR formation zone. In particular, NAA-treated *CASP1_pro_::shy2-2* displayed a larger zone where TCSn::GFP signal was subdued compared to the wild type (Fig. 3B). In the root branching zone, we observed a stronger TCSn::GFP signal in 2,4-D-treated *CASP1_pro_::shy2-2* than Col-0, unlike in IAA- and NAA-treated plants where the TCSn::GFP signals appeared similar between the mutant and wild type (Fig. 3B).

### Endodermal Aux/IAA-mediated signaling is regulates the onset of AUX1:VENUS in the XPP

IAA, NAA and 2,4-D have different transport characteristics in plants. We hypothesized that their different efficiency in rescuing the *CASP1_pro_::shy2-2* LR phenotype could be due to mis-regulation of auxin transporters. To test this, we characterized the expression and localization of the well characterized auxin efflux carrier, PIN-FORMED 1 (PIN1) and importer AUXIN RESISTANT 1 (AUX1) (Bennett *et al*., 1996; Gälweiler *et al*., 1998; Benková *et al*., 2003). Using the *AUX1:VENUS* reporter line (Swarup *et al*., 2004), crossed into *CASP1_pro_::shy2-2*, we observed that the expression of this major auxin influx carrier was almost absent in the LR formation zone and lower compared to the wild type, whereas we observed no differences in *AUX1:VENUS* expression towards the LR branching zone between the two genotypes (Fig. 4A). This correlates with the described expression pattern of the CASP1 promoter that regulates the expression of *shy2-2* to inhibit Aux/IAA-mediated signaling in differentiated endodermal cells (Vermeer *et al*., 2014). Interestingly, treatment with any of the tested auxins was able to recover the *AUX1:VENUS* expression level in *CASP1_pro_::shy2-2* to match the AUX1:VENUS pattern in wild type roots (Fig. 4A), suggesting a possible correlation between the absence of LR phenotype and AUX1 levels in the LR formation zone in *CASP1_pro_::shy2-2*. In contrast, the expression of PIN1, appeared similar to the wild type and did not seem to be influenced by any of the auxin treatments in *CASP1_pro_::shy2-2* (Fig. 4B).

**Figure 4.**
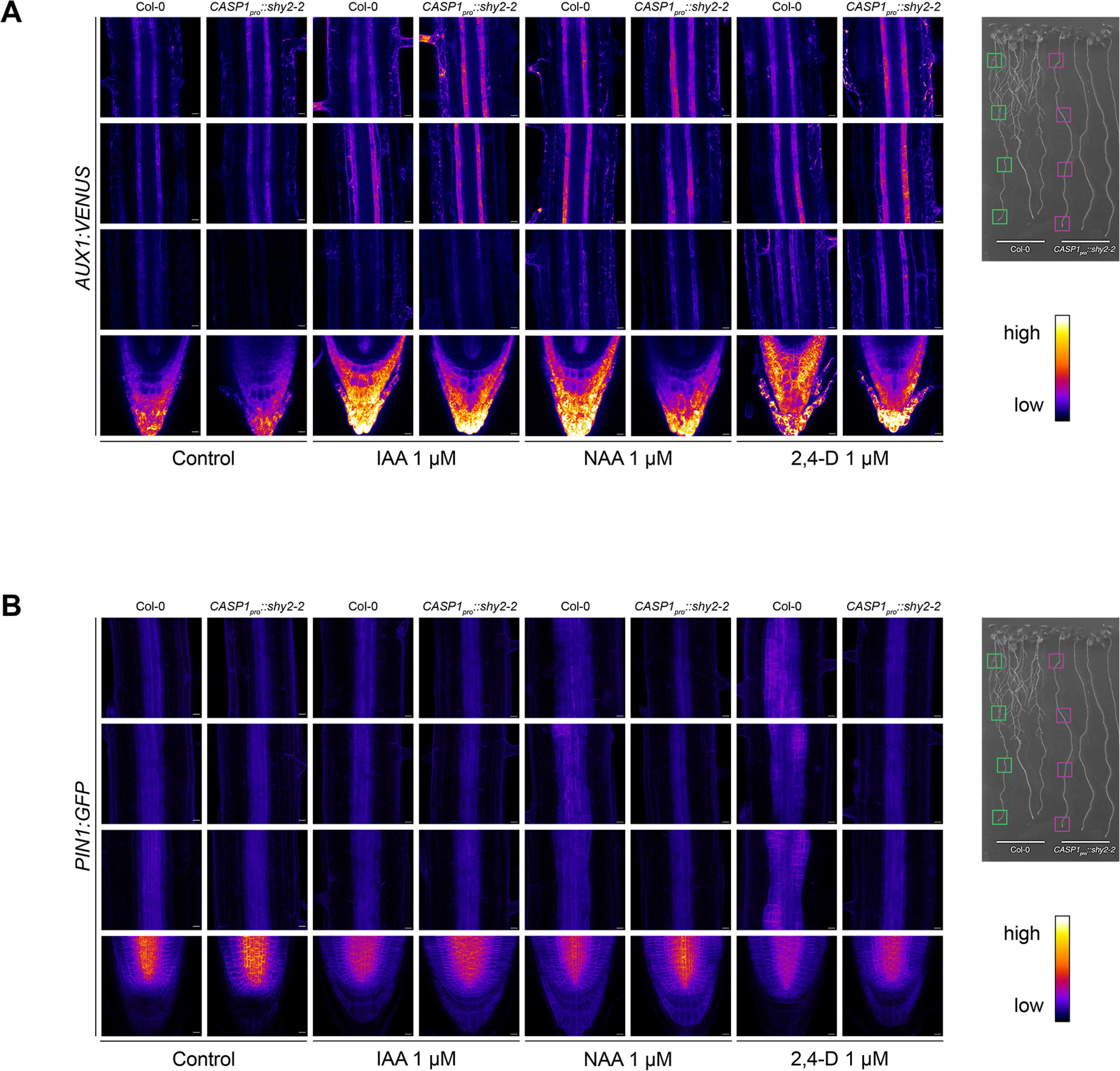
Mis-regulation of auxin transporters in *CASP1_pro_::shy2-2*. 5-day old seedlings of auxin transporter reporter lines, *AUX1:mVENUS* (**A**) and *PIN1:GFP* (**B**), with Col-0 and *CASP1_pro_::shy2-2* backgrounds, treated with IAA, NAA and 2,4-D at 1 μM for 24 hours, were fixed and cleared with ClearSee, and imaged using multiphoton microscope. Relative positions of observation along the primary root are marked with green (Col-0) and magenta (*CASP1_pro_::shy2-2*) boxes. Scale bar = 10 μm; min. 5 images per condition, 3 independent experiments.

**Figure 5.**
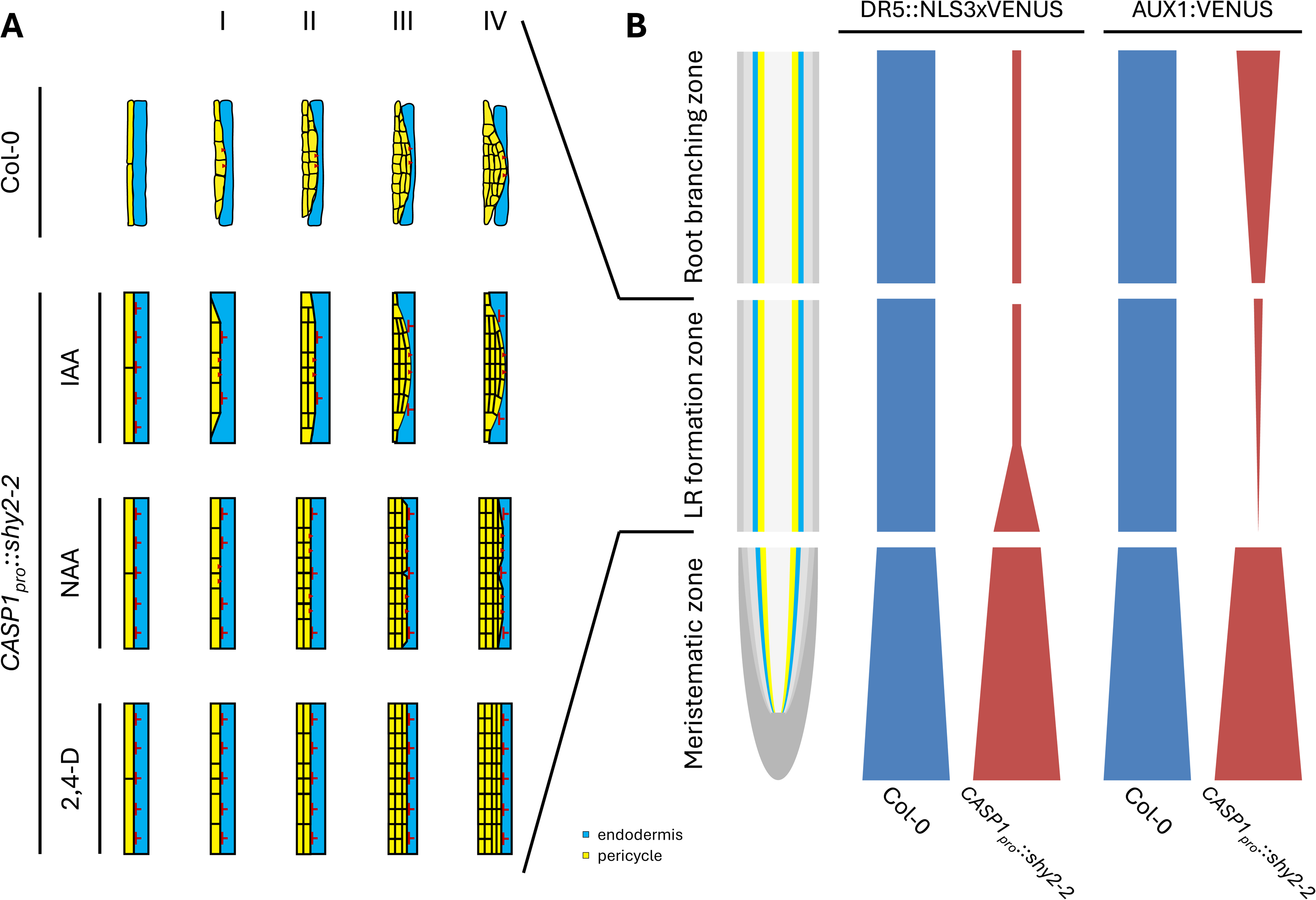
A tightly regulated auxin signaling landscape is required for spatial accommodation of lateral roots in Arabidopsis. **A.** Schematic representation of LRP development up to stage IV in Col-0 (adapted from Stoeckle *et al*., 2018 and auxin-treated *CASP1_pro_::shy2-2*. Red perpendicular lines indicate endodermal mechanical block against LRP outgrowth. Red arrowheads indicate PIN1-mediated auxin flow. **B.** Levels of auxin-induced auxin signaling and AUX1 expression across the root branching zone, the LR formation zone and the meristematic zone in *CASP1_pro_::shy2-2* (red) compared to Col-0 (blue). In auxin-treated *CASP1_pro_::shy2-2*, stronger DR5::NLS3xVENUS signal and weaker AUX1:VENUS signal correlate at the younger end of the LR formation zone.

## DISCUSSION

Several studies have used synthetic, and often more stable, auxins such as NAA and 2,4-D to mimic the effects of IAA due to their comparable affinity in TIR1 binding (Kepinski & Leyser, 2004; Dharmasiri *et al*., 2005; Calderón Villalobos *et al*., 2012; Karami *et al*., 2023). Nevertheless, their effects are not strictly the same, because IAA, NAA and 2,4-D have dissimilar intercellular transport mechanisms (Delbarre *et al*., 1996). IAA can enter the cell partially by diffusion or by active transport via AUXIN RESISTANT (AUX1)/ LIKE-AUX1 (AUX/LAX) influx carriers (Bennett *et al*., 1996; Delbarre *et al*., 1996). NAA can freely diffuse into the cells or be transported through the influx carriers (Delbarre *et al*., 1996). The accumulation level of IAA or NAA in the cell is controlled by the auxin efflux transporters like the PIN-FORMED (PINs) family or the ATP-BINDING CASSETTE sub-family B (ABCBs), whereas 2,4-D is can be imported via the influx carriers, but cannot be exported via the efflux carriers as efficiently (Delbarre *et al*., 1996; Cho & Cho, 2013).

### Different efficacy in LR induction versus emergence by auxin treatment in *CASP1_pro_::shy2-2*

As expected, IAA, being the native auxin in plants, was the most effective in rescuing the LR phenotype in *CASP1_pro_::shy2-2*. IAA induced the formation of LRPs with a morphology that resembled the wild type, albeit that LRP patterning and cell division were not fully preserved (Fig. 1A-B, 2A & 5A). However, it is likely that this could be a result of a potential disturbance of the endogenous auxin gradient that is required for proper LR morphogenesis. NAA, despite sharply increasing LR initiation, did not maintain the standard distributing pattern nor the morphological properties of LRP in *CASP1_pro_::shy2-2* compared to Col-0 (Fig 1A, C, F, I, 2 & S2). 2,4-D, by contrast, severely disturbed LRP morphology as well as LR emergence in *CASP1_pro_::shy2-2* (Fig. 1 & 2A). Thus it appears that the differences in the auxin signaling and intercellular transport pathways of IAA, NAA and 2,4-D are relevant to the responses in *CASP1_pro_::shy2-2* roots observed after their treatments, respectively. Indeed, NAA and 2,4-D, as non-native auxins to plants, could not rescue the LR phenotype in *CASP1_pro_::shy2-2* (Fig. 1). NAA with its high cell-to-cell mobility was able to trigger XPP cell divisions along the primary root and give rise to some dome-shaped LRPs in *CASP1_pro_::shy2-2* roots. In contrast, 2,4-D induced massive cell proliferations in the XPP of *CASP1_pro_::shy2-2* roots compared to wild type and was incapable of inducing dome-shaped LRP. This is most likely due an even more inefficient export of 2,4-D from XPP cells in *CASP1_pro_::shy2-2* roots. As a result, 2,4-D remains “trapped” within XPP cells causing repeated divisions. Moreover, the strong endodermal mechanical resistance in *CASP1_pro_::shy2-2* roots might have favored anticlinal division rather than periclinal, resulting in larger auxin-induced cell proliferation zone and wider LR-like structures (Fig 2A-C). In addition, the observed *CASP1_pro_::shy2-2* phenotypes after NAA and 2,4-D application resemble the phenotype of the *gnom^R5^* mutant treated with the same auxins (Geldner *et al*., 2004). This could imply that the recycling of the PINs or other membrane proteins could be affected in *CASP1_pro_::shy2-2*. Interestingly, a weak allele of *GNOM* has been described in the regulation of the LR founder cell specification in *Arabidopsis* (Wachsman *et al*., 2020). This strong effect could be attributed to the fact that when the 5-day old seedlings were transferred, the *CASP1_pro_::shy2-2* plants already exhibit a non-responsive endodermis. The interrupted endodermis – pericycle communication would likely continue to generate increased endodermal resistance towards the growing LRPs, resulting in the observed problems with LRP morphology in auxin-treated *CASP1_pro_::shy2-2* roots.

### Auxin and cytokinin signaling in *CASP1_pro_::shy2-2*

Being a determinant of cell division, elongation and differentiation, auxin has been widely considered a key signaling molecule involved in the regulation of LR organogenesis. In *Arabidopsis*, overexpression of a stabilized version of *SHY2*, *shy2-2*, in differentiated endodermal cells disrupts local auxin signaling required to regulate LR formation (Vermeer *et al*., 2014). However, SHY2 has already been linked to cytokinin signaling in the root apical meristem (Ioio *et al*., 2008). Because the transduction of auxin and cytokinin signals have counteracting yet complementary effects on LR development (Laplaze *et al*., 2007; Bielach *et al*., 2012; Kieber & Schaller, 2018), the deviating *CASP1_pro_::shy2-2* responses could be partly due to different behaviors in auxin and cytokinin signaling upon each auxin treatment condition. Moreover, the auxin-induced, flattened and widened LRPs prior to traversing the endodermis in *CASP1_pro_::shy2-2* roots (Fig. 2) clearly demonstrates that auxin treatments could partially override the impaired endodermal auxin signaling in *CASP1_pro_::shy2-2*. Although we did not observe major differences in the expression pattern of the cytokinin signaling response marker TCSn::GFP (Fig. 3B), it is plausible that cytokinin signaling contributes to spatial accommodation, as TCSn::GFP was shown to be induced in the overlying endodermis during the later stages of LR development (Zürcher *et al*., 2013). However, in general auxin treatments resulted in an increase in DR5::NLS-3xVENUS signal and a decrease in the TCSn::GFP signal, similarly in both genotypes. Based on our results, we propose that a tightly controlled, auxin transport module acting in the XPP and overlying cell layers is required to accommodate LR development. Treatment with auxins with altered transport properties and stability will disrupt this auxin signaling landscape resulting in altered LR morphogenesis.

### Aux/IAA-mediated signaling in the endodermis is required for timing of AUX1 expression in the XPP

Directional intercellular transport is required to deliver auxin from young aerial tissues where it is biosynthesized to distant targets within a tissue or across the plants (Petrášek & Friml, 2009). This transport is co-handled by a number of integral plasma membrane proteins, most prominently including AUX/LAX influx carriers, PIN efflux carriers and ABCB transporters (Bennett *et al*., 1996; Casimiro *et al*., 2001; Benková *et al*., 2003; Santelia *et al*., 2005; Swarup *et al*., 2008; Péret *et al*., 2009; Petrášek & Friml, 2009; Cho & Cho, 2013). Here we provide evidence that the altered auxin responsiveness of *CASP1_pro_::shy2-2* (Fig. 1 and 2) could be linked to a mis-regulation of auxin transporters in this mutant. We did not observe significant changes in *PIN1* expression in the LR developmental zone between Col-0 and *CASP1_pro_::shy2-2* (Fig. 4B). However, the fact that 2,4- D cannot be transported via auxin efflux carriers efficiently suggests that it is unlikely that the strong cell proliferation in *CASP1_pro_::shy2-2* after 2,4-D treatment is caused by altered expression or activity of the PIN family members (Fig 1A and 2A). Unlike 2,4-D, both IAA and NAA are normally exported from the cell via PINs, thus the PIN1-mediated polar transport of these auxins might have remained unchanged in treated *CASP1_pro_::shy2-2.* This could direct cell division up against the endodermis, so LRPs and LRs could be formed consequently (Fig. 1A, 2A & 5A). Interestingly, we showed that *AUX1:VENUS* was repressed in the LR formation zone of *CASP1_pro_::shy2-2* roots, and that auxin treatment could restore this expression (Fig. 4A & 5B). This suggests that an early signal, dependent on Aux/IAA-mediated signaling in the endodermis is required to potentiate the XPP to accumulate AUX1 to initiate the LR developmental program. It would be tempting to speculate that this could be a mechanical signal as it was recently shown that re-organization of the cortical microtubule cytoskeleton in the overlying endodermis is required to accommodate LR initiation (Stoeckle *et al*., 2022).

## AUTHOR CONTRIBUTIONS

TXB and VS performed auxin treatment assays. TXB performed analysis of LR morphogenesis, responses marker dynamics and localization of auxin transporters. SMM provided new reagents. KB and JEMV supervised the project. TXB, KB, and JEMV wrote the manuscript.

## ACKNOWLEDGEMENTS

We would like to thank Prof. Malcolm Bennett (University of Nottingham) for the AUX1 reporter lines. Many thanks to the Vermeer lab members for their continuous feedback during this project.

## DATA AVAILABILITY STATEMENTS

The data that support the findings of this study are available from the corresponding author upon reasonable request.

## SUPPORTING INFORMATION

**Figure S1. Different paths of cellular transport of IAA, NAA and 2,4-D.**

**A**. IAA can be transported into and out of the cells via auxin carriers, or via less efficient passive diffusion. **B.** NAA can be transported into and out of the cells via auxin carriers and passive diffusion. **C.** 2,4-D can only enter the cells via auxin influx carriers, but cannot diffuse nor be efficiently exported via auxin efflux carriers. Solid arrows: main paths of auxin transport; dashed arrows: less efficient paths of auxin transport. (Schematic was inspired by Delbarre *et al*., 1996)

**Figure S2. LR emergence is not fully rescued in auxin-treated *CASP1_pro_::shy2-2*.**

**A**-**C**. Proportion of non-emerged LRs (stage I - VII) in Col-0 (dark green) and *CASP1_pro_::shy2-2* (dark yellow) and emerged LRs (stage VIII) in Col-0 (light green) and *CASP1_pro_::shy2-2* (light yellow), induced by IAA (**A**), NAA (**B**) and 2,4-D (**C**). Different letters indicate Pearson’s χ^2^ test significant difference (p-value < 0.05). **D.** Observed distribution of emerged LRs in correspondence with the root branching zone, the LR formation zone and the meristematic zone in *CASP1_pro_::shy2-2* after extended auxin treatments (red) compared to Col-0 (blue). 5-day old seedlings of Col-0 and *CASP1_pro_::shy2-2* were treated with IAA, NAA and 2,4-D at 0.1, 1 or 10 μM for up to 72 hours. Root phenotyping was performed on no less than 18 plants per condition in 3 replicates.

Figures are submitted as separate documents.

## REFERENCES

Benková, E., Michniewicz, M., Sauer, M., Teichmann, T., Seifertová, D., Jürgens, G., Friml, J., 2003. Local, Efflux-Dependent Auxin Gradients as a Common Module for Plant Organ Formation. Cell 115, 591–602. 10.1016/S0092-8674(03)00924-3

Bennett, M.J., Marchant, A., Green, H.G., May, S.T., Ward, S.P., Millner, P.A., Walker, A.R., Schulz, B., Feldmann, K.A., 1996. *Arabidopsis* AUX1 Gene: A Permease-Like Regulator of Root Gravitropism. Science 273, 948–950. 10.1126/science.273.5277.948

Bielach, A., Podlešáková, K., Marhavý, P., Duclercq, J., Cuesta, C., Müller, B., Grunewald, W., Tarkowski, P., Benková, E., 2012. Spatiotemporal Regulation of Lateral Root Organogenesis in *Arabidopsis* by Cytokinin. Plant Cell 24, 3967–3981. 10.1105/tpc.112.103044

Calderón Villalobos, L.I.A., Lee, S., De Oliveira, C., Ivetac, A., Brandt, W., Armitage, L., Sheard, L.B., Tan, X., Parry, G., Mao, H., Zheng, N., Napier, R., Kepinski, S., Estelle, M., 2012. A combinatorial TIR1/AFB–Aux/IAA co-receptor system for differential sensing of auxin. Nat. Chem. Biol. 8, 477–485. 10.1038/nchembio.926

Cancé, C., Martin-Arevalillo, R., Boubekeur, K., Dumas, R., 2022. Auxin response factors are keys to the many auxin doors. New Phytol. 235, 402–419. 10.1111/nph.18159

Casimiro, I., Marchant, A., Bhalerao, R.P., Beeckman, T., Dhooge, S., Swarup, R., Graham, N., Inzé, D., Sandberg, G., Casero, P.J., Bennett, M., 2001. Auxin Transport Promotes *Arabidopsis* Lateral Root Initiation. Plant Cell 13, 843–852. 10.1105/tpc.13.4.843

Cho, M., Cho, H., 2013. The function of ABCB transporters in auxin transport. Plant Signal. Behav. 8, e22990. 10.4161/psb.22990

de Jesus Vieira Teixeira, C., Bellande, K., van der Schuren, A., O’Connor, D., Hardtke, C.S., Vermeer, J.E.M., 2024. An atlas of Brachypodium distachyon lateral root development. Biol. Open 13, bio060531. 10.1242/bio.060531

De Rybel, B., Vassileva, V., Parizot, B., Demeulenaere, M., Grunewald, W., Audenaert, D., Van Campenhout, J., Overvoorde, P., Jansen, L., Vanneste, S., Möller, B., Wilson, M., Holman, T., Van Isterdael, G., Brunoud, G., Vuylsteke, M., Vernoux, T., De Veylder, L., Inzé, D., Weijers, D., Bennett, M.J., Beeckman, T., 2010. A Novel Aux/IAA28 Signaling Cascade Activates GATA23-Dependent Specification of Lateral Root Founder Cell Identity. Curr. Biol. 20, 1697–1706. 10.1016/j.cub.2010.09.007

De Smet, I., Lau, S., Voß, U., Vanneste, S., Benjamins, R., Rademacher, E.H., Schlereth, A., De Rybel, B., Vassileva, V., Grunewald, W., Naudts, M., Levesque, M.P., Ehrismann, J.S., Inzé, D., Luschnig, C., Benfey, P.N., Weijers, D., Van Montagu, M.C.E., Bennett, M.J., Jürgens, G., Beeckman, T., 2010. Bimodular auxin response controls organogenesis in *Arabidopsis*. Proc. Natl. Acad. Sci. 107, 2705–2710. 10.1073/pnas.0915001107

De Smet, I., Tetsumura, T., De Rybel, B., Frey, N.F. dit, Laplaze, L., Casimiro, I., Swarup, R., Naudts, M., Vanneste, S., Audenaert, D., Inzé, D., Bennett, M.J., Beeckman, T., 2007. Auxin-dependent regulation of lateral root positioning in the basal meristem of *Arabidopsis*. Development 134, 681–690. 10.1242/dev.02753

Delbarre, A., Muller, P., Imhoff, V., Guern, J., 1996. Comparison of mechanisms controlling uptake and accumulation of 2,4-dichlorophenoxy acetic acid, naphthalene-1-acetic acid, and indole-3-acetic acid in suspension-cultured tobacco cells. Planta 198, 532–541. 10.1007/BF00262639

Dharmasiri, N., Dharmasiri, S., Estelle, M., 2005. The F-box protein TIR1 is an auxin receptor. Nature 435, 441–445. 10.1038/nature03543

Du, Y., Scheres, B., 2018. Lateral root formation and the multiple roles of auxin. J. Exp. Bot. 69, 155–167. 10.1093/jxb/erx223

Gälweiler, L., Guan, C., Müller, A., Wisman, E., Mendgen, K., Yephremov, A., Palme, K., 1998. Regulation of Polar Auxin Transport by AtPIN1 in *Arabidopsis* Vascular Tissue. Science 282, 2226–2230. 10.1126/science.282.5397.2226

Geldner, N., Dénervaud-Tendon, V., Hyman, D.L., Mayer, U., Stierhof, Y.-D., Chory, J., 2009. Rapid, combinatorial analysis of membrane compartments in intact plants with a multicolor marker set. Plant J. 59, 169–178. 10.1111/j.1365-313X.2009.03851.x

Geldner, N., Richter, S., Vieten, A., Marquardt, S., Torres-Ruiz, R.A., Mayer, U., Jürgens, G., 2004. Partial loss-of-function alleles reveal a role for GNOM in auxin transport-related, post-embryonic development of *Arabidopsis*. Development 131, 389–400. 10.1242/dev.00926

Gifford, M.L., Xu, G., Dupuy, L.X., Vissenberg, K., Rebetzke, G., 2024. Root architecture and rhizosphere–microbe interactions. J. Exp. Bot. 75, 503. 10.1093/jxb/erad488

Goh, T., Kasahara, H., Mimura, T., Kamiya, Y., Fukaki, H., 2012. Multiple AUX/IAA–ARF modules regulate lateral root formation: the role of *Arabidopsis* SHY2/IAA3-mediated auxin signalling. Philos. Trans. R. Soc. B Biol. Sci. 367, 1461–1468. 10.1098/rstb.2011.0232

Gray, W.M., Muskett, P.R., Chuang, H., Parker, J.E., 2003. *Arabidopsis* SGT1b Is Required for SCFTIR1-Mediated Auxin Response. Plant Cell 15, 1310–1319. 10.1105/tpc.010884

Guilfoyle, T.J., Hagen, G., 2007. Auxin response factors. *Curr. Opin. Plant Biol.*, Cell Signalling and Gene Regulation 10, 453–460. 10.1016/j.pbi.2007.08.014

Heisler, M.G., Ohno, C., Das, P., Sieber, P., Reddy, G.V., Long, J.A., Meyerowitz, E.M., 2005. Patterns of Auxin Transport and Gene Expression during Primordium Development Revealed by Live Imaging of the *Arabidopsis* Inflorescence Meristem. Curr. Biol. 15, 1899–1911. 10.1016/j.cub.2005.09.052

Ioio, R.D., Nakamura, K., Moubayidin, L., Perilli, S., Taniguchi, M., Morita, M.T., Aoyama, T., Costantino, P., Sabatini, S., 2008. A Genetic Framework for the Control of Cell Division and Differentiation in the Root Meristem. Science 322, 1380–1384. 10.1126/science.1164147

Karami, O., de Jong, H., Somovilla, V.J., Villanueva Acosta, B., Sugiarta, A.B., Ham, M., Khadem, A., Wennekes, T., Offringa, R., 2023. Structure–activity relationship of 2,4-D correlates auxinic activity with the induction of somatic embryogenesis in *Arabidopsis thaliana*. Plant J. 116, 1355–1369. 10.1111/tpj.16430

Kepinski, S., Leyser, O., 2004. Auxin-induced SCFTIR1–Aux/IAA interaction involves stable modification of the SCFTIR1 complex. Proc. Natl. Acad. Sci. 101, 12381–12386. 10.1073/pnas.0402868101

Kieber, J.J., Schaller, G.E., 2018. Cytokinin signaling in plant development. Development 145, dev149344. 10.1242/dev.149344

Kurihara, D., Mizuta, Y., Sato, Y., Higashiyama, T., 2015. ClearSee: a rapid optical clearing reagent for whole-plant fluorescence imaging. Development 142, 4168–4179. 10.1242/dev.127613

Laplaze, L., Benkova, E., Casimiro, I., Maes, L., Vanneste, S., Swarup, R., Weijers, D., Calvo, V., Parizot, B., Herrera-Rodriguez, M.B., Offringa, R., Graham, N., Doumas, P., Friml, J., Bogusz, D., Beeckman, T., Bennett, M., 2007. Cytokinins Act Directly on Lateral Root Founder Cells to Inhibit Root Initiation. Plant Cell 19, 3889–3900. 10.1105/tpc.107.055863

Lavenus, J., Goh, T., Roberts, I., Guyomarc’h, S., Lucas, M., De Smet, I., Fukaki, H., Beeckman, T., Bennett, M., Laplaze, L., 2013. Lateral root development in *Arabidopsis*: fifty shades of auxin. Trends Plant Sci. 18, 450–458. 10.1016/j.tplants.2013.04.006

Marhavý, P., Duclercq, J., Weller, B., Feraru, E., Bielach, A., Offringa, R., Friml, J., Schwechheimer, C., Murphy, A., Benková, E., 2014. Cytokinin Controls Polarity of PIN1- Dependent Auxin Transport during Lateral Root Organogenesis. Curr. Biol. 24, 1031– 1037. 10.1016/j.cub.2014.04.002

Moreno-Risueno, M.A., Van Norman, J.M., Moreno, A., Zhang, J., Ahnert, S.E., Benfey, P.N., 2010. Oscillating Gene Expression Determines Competence for Periodic *Arabidopsis* Root Branching. Science 329, 1306–1311. 10.1126/science.1191937

Nenadić, M., Vermeer, J.E.M., 2021. Dynamic cytokinin signalling landscapes during lateral root formation in *Arabidopsis*. Quant. Plant Biol. 2, e13. 10.1017/qpb.2021.13

Péret, B., De Rybel, B., Casimiro, I., Benková, E., Swarup, R., Laplaze, L., Beeckman, T., Bennett, M.J., 2009. *Arabidopsis* lateral root development: an emerging story. Trends Plant Sci. 14, 399–408. 10.1016/j.tplants.2009.05.002

Perotti, M.F., Ariel, F.D., Chan, R.L., 2020. Lateral root development differs between main and secondary roots and depends on the ecotype. Plant Signal. Behav. 15, 1755504. 10.1080/15592324.2020.1755504

Petrášek, J., Friml, J., 2009. Auxin transport routes in plant development. Development 136, 2675– 2688. 10.1242/dev.030353

Santelia, D., Vincenzetti, V., Azzarello, E., Bovet, L., Fukao, Y., Düchtig, P., Mancuso, S., Martinoia, E., Geisler, M., 2005. MDR-like ABC transporter AtPGP4 is involved in auxin- mediated lateral root and root hair development. FEBS Lett. 579, 5399–5406. 10.1016/j.febslet.2005.08.061

Santos Teixeira, J.A., ten Tusscher, K.H., 2019. The Systems Biology of Lateral Root Formation: Connecting the Dots. *Mol. Plant*, Plant Systems Biology 12, 784–803. 10.1016/j.molp.2019.03.015

Schäfer, E.D., Owen, M.R., Band, L.R., Farcot, E., Bennett, M.J., Lynch, J.P., 2022. Modeling root loss reveals impacts on nutrient uptake and crop development. Plant Physiol. 190, 2260– 2278. 10.1093/plphys/kiac405

Schindelin, J., Arganda-Carreras, I., Frise, E., Kaynig, V., Longair, M., Pietzsch, T., Preibisch, S., Rueden, C., Saalfeld, S., Schmid, B., Tinevez, J.-Y., White, D.J., Hartenstein, V., Eliceiri, K., Tomancak, P., Cardona, A., 2012. Fiji: an open-source platform for biological-image analysis. Nat. Methods 676–682. doi:10.1038/nmeth.2019

Stoeckle, D., Reyes-Hernández, B.J., Barro, A.V., Nenadić, M., Winter, Z., Marc-Martin, S., Bald, L., Ursache, R., Fujita, S., Maizel, A., Vermeer, J.E., 2022. Microtubule-based perception of mechanical conflicts controls plant organ morphogenesis. Sci. Adv. 8, eabm4974. 10.1126/sciadv.abm4974

Stoeckle, D., Thellmann, M., Vermeer, J.E., 2018. Breakout — lateral root emergence in *Arabidopsis thaliana*. Curr. Opin. Plant Biol. 41, 67–72. 10.1016/j.pbi.2017.09.005

Swarup, K., Benková, E., Swarup, R., Casimiro, I., Péret, B., Yang, Y., Parry, G., Nielsen, E., De Smet, I., Vanneste, S., Levesque, M.P., Carrier, D., James, N., Calvo, V., Ljung, K., Kramer, E., Roberts, R., Graham, N., Marillonnet, S., Patel, K., Jones, J.D.G., Taylor, C.G., Schachtman, D.P., May, S., Sandberg, G., Benfey, P., Friml, J., Kerr, I., Beeckman, T., Laplaze, L., Bennett, M.J., 2008. The auxin influx carrier LAX3 promotes lateral root emergence. Nat. Cell Biol. 10, 946–954. 10.1038/ncb1754

Swarup, R., Kargul, J., Marchant, A., Zadik, D., Rahman, A., Mills, R., Yemm, A., May, S., Williams, L., Millner, P., Tsurumi, S., Moore, I., Napier, R., Kerr, I.D., Bennett, M.J., 2004. Structure-Function Analysis of the Presumptive *Arabidopsis* Auxin Permease AUX1[W]. Plant Cell 16, 3069–3083. 10.1105/tpc.104.024737

Vanneste, S., De Rybel, B., Beemster, G.T.S., Ljung, K., De Smet, I., Van Isterdael, G., Naudts, M., Iida, R., Gruissem, W., Tasaka, M., Inzé, D., Fukaki, H., Beeckman, T., 2005. Cell Cycle Progression in the Pericycle Is Not Sufficient for SOLITARY ROOT/IAA14- Mediated Lateral Root Initiation in *Arabidopsis thaliana*. Plant Cell 17, 3035–3050. 10.1105/tpc.105.035493

Vermeer, J.E.M., Geldner, N., 2015. Lateral root initiation in *Arabidopsis thaliana*: a force awakens. F1000Prime Rep. 7. 10.12703/P7-32

Vermeer, J.E.M., von Wangenheim, D., Barberon, M., Lee, Y., Stelzer, E.H.K., Maizel, A., Geldner, N., 2014. A Spatial Accommodation by Neighboring Cells Is Required for Organ Initiation in *Arabidopsis*. Science 343, 178–183. 10.1126/science.1245871

Vilches Barro, A., Stöckle, D., Thellmann, M., Ruiz-Duarte, P., Bald, L., Louveaux, M., von Born, P., Denninger, P., Goh, T., Fukaki, H., Vermeer, J.E.M., Maizel, A., 2019. Cytoskeleton Dynamics Are Necessary for Early Events of Lateral Root Initiation in *Arabidopsis*. Curr. Biol. CB 29, 2443–2454.e5. 10.1016/j.cub.2019.06.039

Wachsman, G., Zhang, J., Moreno-Risueno, M.A., Anderson, C.T., Benfey, P.N., 2020. Cell wall remodeling and vesicle trafficking mediate the root clock in *Arabidopsis*. Science 370, 819–823. 10.1126/science.abb7250

Zenser, N., Ellsmore, A., Leasure, C., Callis, J., 2001. Auxin modulates the degradation rate of Aux/IAA proteins. Proc. Natl. Acad. Sci. 98, 11795–11800. 10.1073/pnas.211312798

Zürcher, E., Tavor-Deslex, D., Lituiev, D., Enkerli, K., Tarr, P.T., Müller, B., 2013. A Robust and Sensitive Synthetic Sensor to Monitor the Transcriptional Output of the Cytokinin Signaling Network in Planta. Plant Physiol. 161, 1066–1075. 10.1104/pp.112.211763

