## Supplementary figures and images for "A tightly regulated auxin signaling landscape is required for spatial accommodation of lateral roots in *Arabidopsis*"

### Supplemental Figure S1

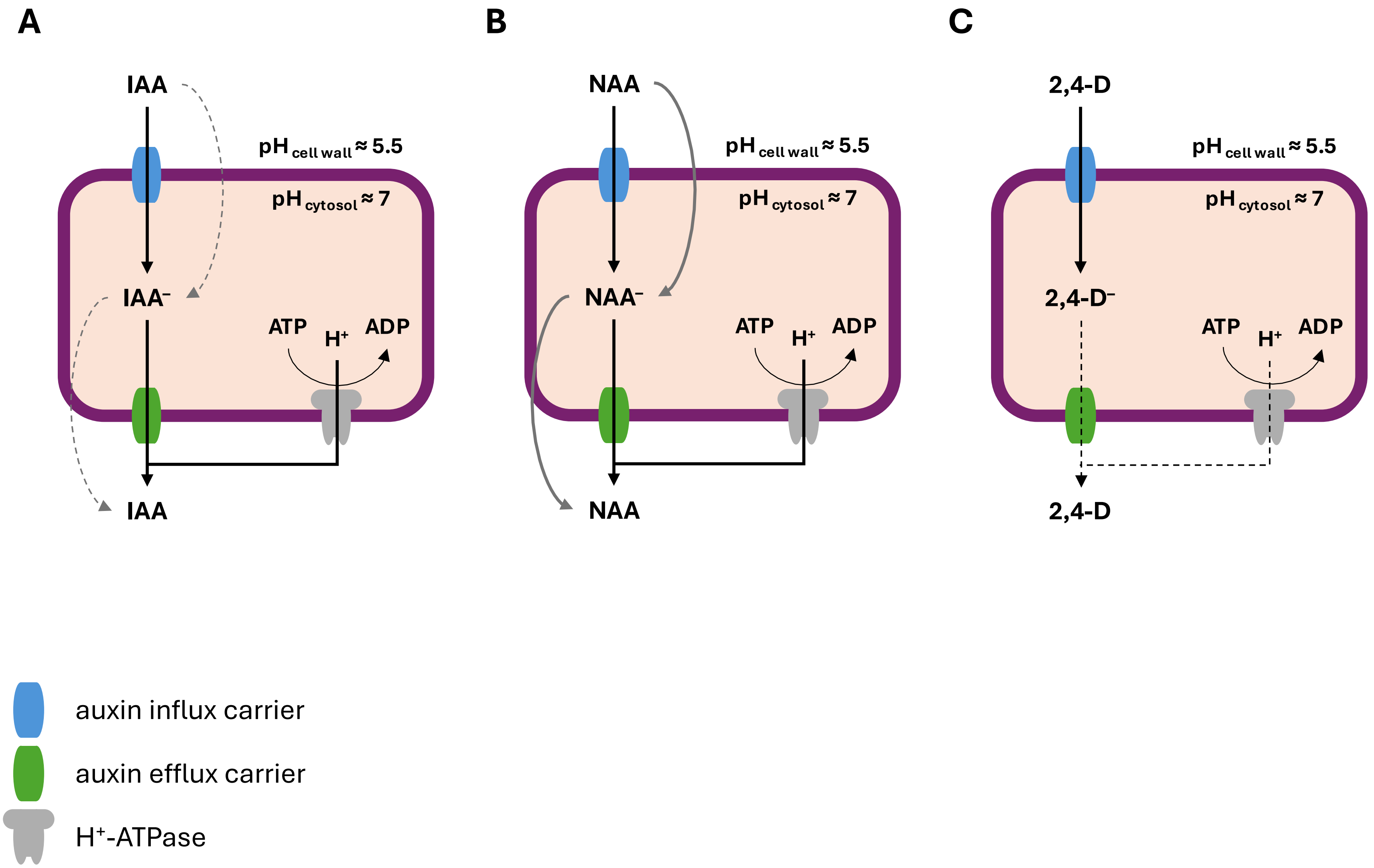

### Supplemental Figure S2

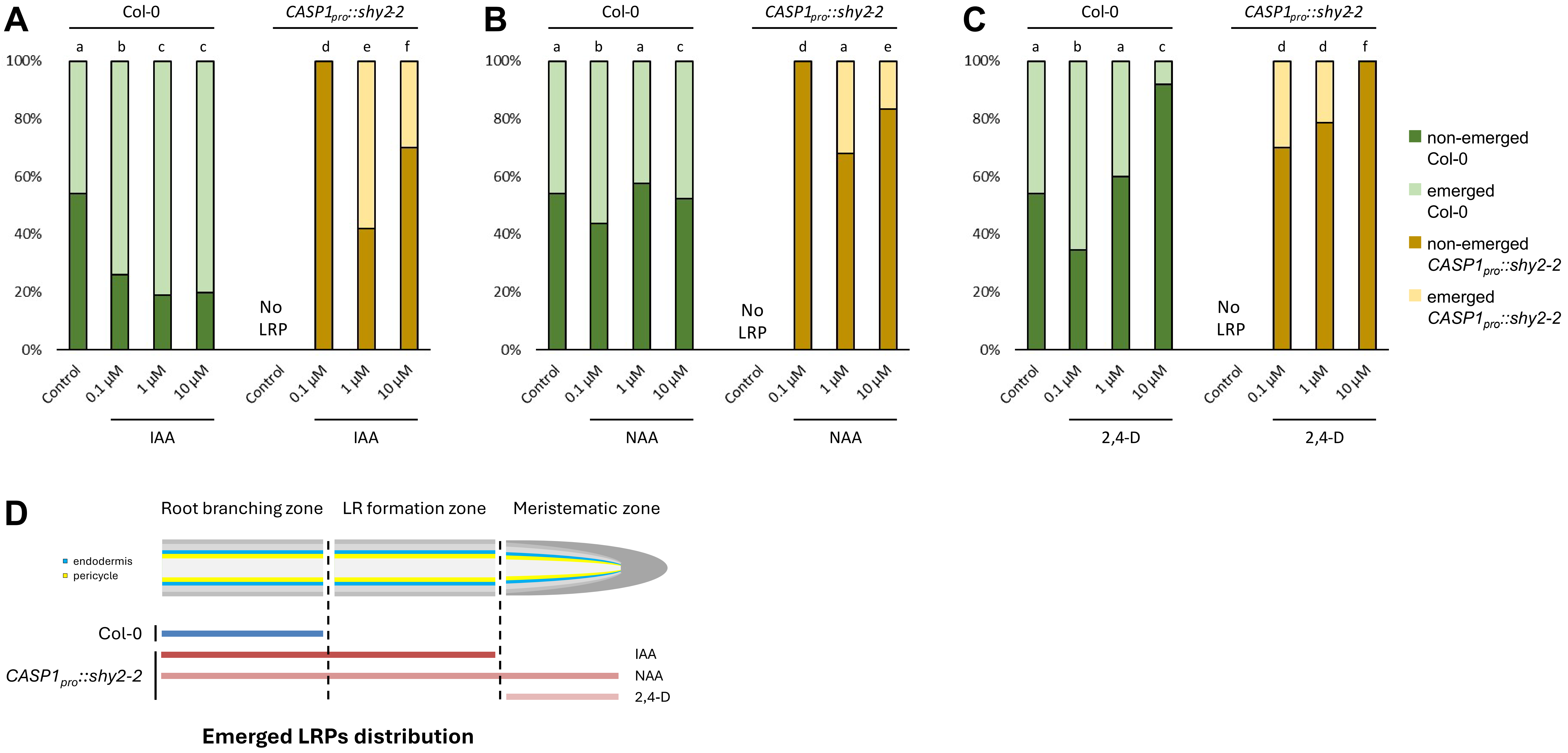
